# Superoxide Dismutases maintain niche homeostasis in stem cell populations

**DOI:** 10.1101/2023.12.06.570086

**Authors:** Olivia Majhi, Aishwarya Chhatre, Tanvi Chaudhary, Devanjan Sinha

**Author notes:** To whom correspondence should be addressed: **Address for correspondence:** Dr. Devanjan Sinha, Department of Zoology, Institute of Science, Banaras Hindu University, Varanasi – 221005, Uttar Pradesh, India, India.

## Abstract

Reactive oxygen species (ROS), predominantly derived from mitochondrial respiratory complexes, have emerged as key molecules influencing cell fate decisions like maintenance and differentiation. These redox-dependent events are mainly considered to be cell intrinsic in nature, on the contrary our observations indicate involvement of these oxygen-derived entities as intercellular communicating agents. In *Drosophila* male germline, Germline Stem Cells (GSCs) and neighbouring Cyst Stem Cells (CySCs) maintain differential redox thresholds where CySC have higher redox-state compared to the adjacent GSCs. Disruption of the redox equilibrium between the two adjoining stem cell populations by depleting Superoxide Dismutases (SODs) especially Sod1 results in deregulated niche architecture and loss of GSCs, which was mainly attributed to loss of contact-based receptions and uncontrolled CySC proliferation due to ROS-mediated activation of self-renewing signals. Our observations hint towards the crucial role of differential redox states where CySCs containing higher ROS function not only as a source of their own maintenance cues but also serve as non-autonomous redox moderators of GSCs. Our findings underscore the complexity of niche homeostasis and predicate the importance of intercellular redox communication in understanding stem cell microenvironments.

## Introduction

Studies from the past few decades have shown the apparent role of ROS in influencing various biological processes^1–3^. ROS are usually produced in specific cellular compartments close to their target molecules to regulate important signaling pathways.^4^. Mitochondria, a major source of oxidant species^5,6^, localize dynamically towards nucleus, effecting oxidation induced reshaping of gene expression profiles^7^. Localized ROS production are contributed by NADH oxidases distributed across different cellular regions^8^. Intracellular hydrogen peroxide gradients, maintained by thioredoxin system, allow intercompartmental exchanges among endoplasmic reticulum (ER), mitochondria and peroxisomes^9–11^.

The process of redox relays is conserved and plays a fundamental role in self-renewal and differentiation of stem cell populations^12^. Stem cells whether embryonic or adult generally maintain a low redox profile, barring few exceptions and are characterized by subdued mitochondrial respiration^13–17^. Levels of ROS are tightly regulated and elevated amounts promote early differentiation and atypical stem cell behaviour^18^. However, leading evidences suggest that many transcription factors require oxidative environment for the maintenance of pluripotent states. For instance, physiological ROS levels play a crucial role in genome maintenance of embryonic stem cells^13^. Multipotent hematopoietic progenitors, intestinal stem cells ^19^ and neural stem cells require relatively high baseline redox for their maintenance^14,16,20^. Suppression of Nox system or mitochondrial ROS compromises the maintenance of these self-renewing populations and promote their differentiation or death^18^. These evidences point towards the essentiality of a well-tuned redox state for balancing the pluripotent and differentiated states. However, how stem cells regulate their redox potential by possessing a restrained oxidant system is not very clear.

We addressed this fundamental question in two-stem cell population-based niche architecture in *Drosophila* testis. The testicular stem cell niche is composed of a central cluster of somatic cells called the hub which contacts eight to eleven GSCs arranged in a round array^21,22^. A pair of cyst stem cells (CySCs) enclose each GSC and make their independent connections with the hub and GSCs via adherens junctions^23–28^ The hub cells secrete self-renewal factors essential for both GSCs and CySCs maintenance involving signalling cascades like Jak-Stat signalling^29,30^, BMP (Bone Morphogenetic Signalling)^31^, Hedgehog signalling^32^. The hub and CySCs produce BMPs which repress GSC differentiation by suppressing transcription of bag-of-marbles (Bam)^31,33^. Asymmetric division of both GSC and CySC is essential for proper cyst formation^34,35^. A developing cyst contains a dividing and differentiating gonialblast encircled by cyst cells that provide nourishment to developing spermatids^36,37^. However, the maintenance of GSCs in the spatially controlled microenvironment is still not very clear.

We found the existence of a balanced differential redox state between GSC and CySC to be essential for niche homeostasis. CySCs by virtue of their higher redox threshold and more clustered mitochondria generated an intercellular redox state that affected the physiological ROS levels in GSCs. Alterations in CySC redox state affected the self-renewal and differentiation propensities of the GSCs. Intercellular redox imbalance disrupts niche homeostasis by inducing premature differentiation of germline stem cells (GSCs) and aberrant proliferation of cyst stem cells (CySCs), driven by activation of pro-proliferative signaling pathways and attenuation of cell–cell contact–mediated communication. Our results indicated a sophisticated interplay between these two stem cell populations where a higher redox state in CySCs not only supports their self-renewing processes but also non-autonomously promotes the maintenance of GSCs.

## Results

### CySCs maintain higher differential ROS in comparison to adjacent GSCs

The apical region of the *Drosophila* testis incorporates a cluster of differentiated cells, the hub which is surrounded by GSCs in a rosette arrangement with each GSC being enclosed by two CySCs (Fig. 1A). Immunostaining the ATP5A subunit of mitochondrial ATPase and marking cell boundary by discs-large (Dlg) indicated different mitochondrial distribution among these two stem cell populations. In wild type adult testis, we observed sparsely populated mitochondria in Vasa^+^ GSCs as compared to their dense distribution in CySCs (Fig. 1B#, C, D, and S1B-B’’). The number of mitochondria per cell was higher in CySC (Fig. 1F). The ATP5A labelling overlapped with *TFAM-GFP*, a mitochondrial transcription factor^38^ that labelled the mitochondria, further confirming the patterning observed between these two stem cell populations (Fig. S1A). Since mitochondria are known to be one of the major producers of ROS in cells, we tested for the relative redox profiles in these two stem cell populations. Mitochondrial dispersion pattern in GSCs and CySCs, corresponded with the intensity variance of the ROS reporter line *gstD1-GFP* ^39^ among the different cell populations at the niche. CySCs (outlined by a dotted boundary) exhibited distinctly higher *gstD1* reporter intensity than the neighbouring GSC (shown with yellow arrowhead) (Fig. 1K and S1C’’’). Quantification of this intensity difference indicated that CySCs exhibited a higher baseline level of ROS compared to GSCs (Fig.1Q). The hub zone (denoted by asterisk) also presented higher *gstD1-GFP* intensity when compared to the surrounding germline (Fig. 1K and S1C’’’).

**Figure 1.**
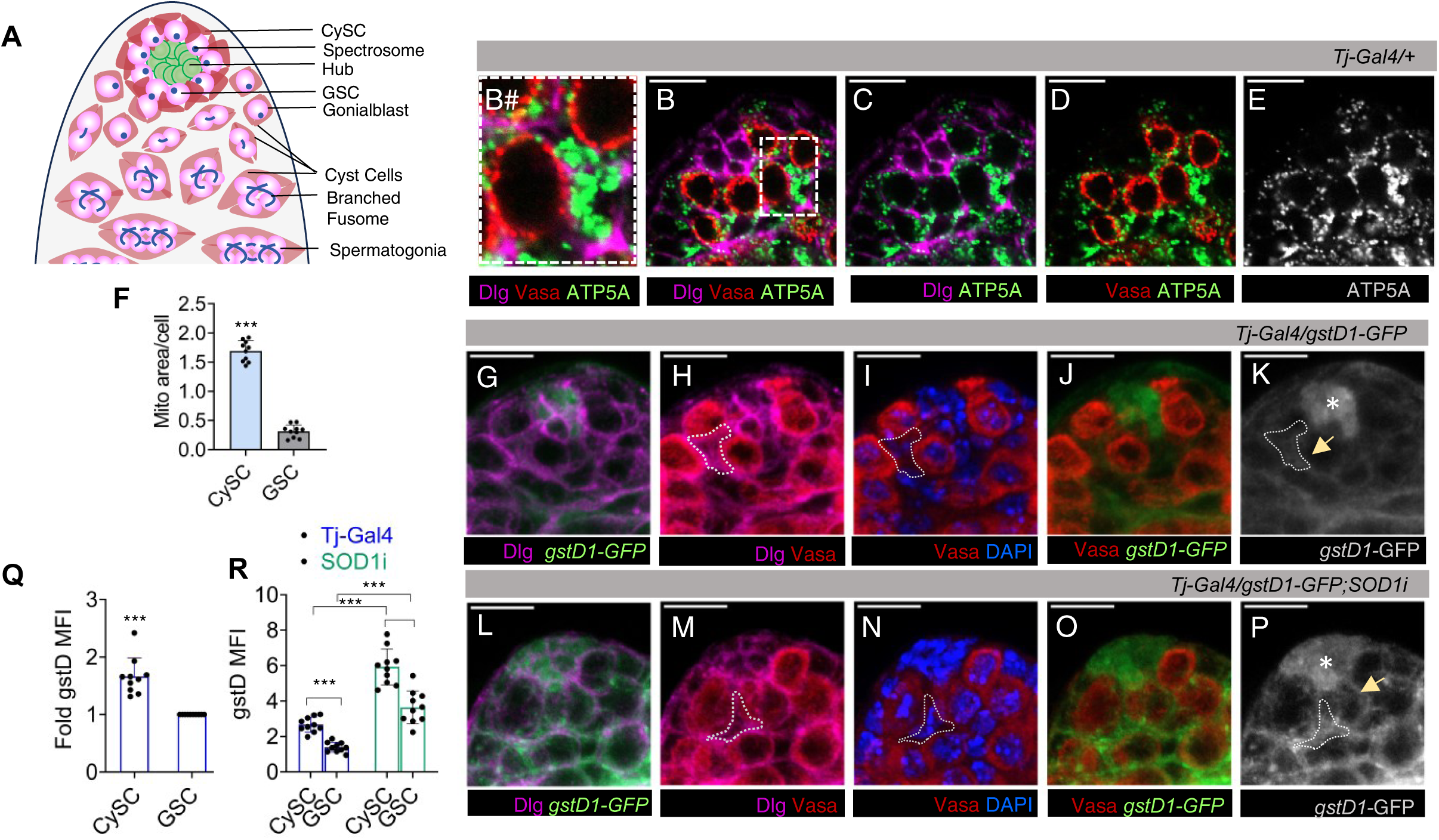
Mitochondrial Distribution and Redox Profile of GSCs and CySCs in Adult *Drosophila* Testes. (A) Schematic representation of adult *Drosophila* testicular niche showing arrangement of different stem cell populations (GSC – Germinal Stem Cell, CySC – Cyst Stem Cell). (B-E) Differential distribution of mitochondria labelled with ATP5A (monochrome/green) in Vasa^+^ GSCs of wild-type fly testis with cellular boundaries marked by Dlg; B# shows digitally zoomed image of the dotted area of B. (F) Quantification of mitochondrial area per cell. Bars represent mean ± s.e.m., n = 10 fields of view; *** *P (unpaired t-test)<0.0001.* (G-P) Redox profiling of testicular stem cell niche using *gstD1* -GFP as intrinsic ROS reporter in control (G-K) and *Tj-Gal4* driven *Sod1RNAi (Sod1i)* testis (L-P). Dotted area denotes the region of CySC occupancy and asterisk denotes the hub. (Q) Quantification of *gstD1-*GFP mean fluorescence intensity (MFI) represented as fold change between GSCs and CySCs in controls. Data denotes mean ± s.e.m,, n = 25, *** *P(unpaired t-test)<0.0001.* (R) Quantification of cell-specific ROS content upon *Sod1RNAi*. Data denotes mean ± s.e.m, n = 25, *** *P(unpaired t-test)<0.0001.* Controls are the indicated driver line crossed with *Oregon R^+^*. Scale bar - 10 µm. See also Figure S1.

Given the ability of free-radicals to diffuse, we hypothesized that the somatically derived cells in the niche, by virtue to their higher ROS state might be maintaining the redox environment of their surroundings. In this study, we evaluated the redox interplay between the two stem cell population in the niche. To test the role of CySC in influencing GSC redox state, we asked if disrupting the superoxide dismutases in CySCs would influence the redox state of GSCs. To deplete SOD1, we used *Tj-Gal4*^40^ driver (*Tj>Sod1i*) that expresses majorly in CySCs and early differentiating cyst cells (CCs) with minor levels of expression in the hub. Depletion of SOD1 majorly in CySCs resulted in a net increase in *gstD1* -GFP intensity in the niche zone (Fig. 1P, R and S1E), and an overall rise in superoxide levels, confirmed through DHE fluorescence analysis (Fig.S1F). Increased *gstD1* labelling due to SOD1 depletion in CySC (denoted by dotted area) (Fig. 1P, S1D’’’) adjacent to Vasa+ GSC (shown with yellow arrow), resulted in concomitant increase in *gstD1* intensity at the Vasa+ GSC (Fig. 1P, S1D’’’). This indicates that the elevated redox state in CySCs led to a corresponding increase in ROS levels in adjacent GSCs (Fig. 1R) suggesting a potential redox crosstalk between these stem cell populations, where the redox state of CySCs might be influencing the oxidative status of neighbouring GSCs. This crosstalk was further validated by subsequent phenotypic analysis.

### Balanced CySC redox profile is crucial for its proliferation and GSC maintenance

Alongside the differential redox profile, we observed that elevated ROS levels in *Tj>Sod1i* had a striking effect on the increase of DAPI^+^ nuclei at the testis tip (Fig. 2A’’). This overcrowding was attributed to both an increase in the number of Tj^+^ CySCs and early differentiating CCs, as well as their positional shift (Fig. 2A’, B’, and E). To further validate these findings, we used an alternative driver line, *C-587-Gal4*, which specifically drives expression in CySCs and CCs. This also resulted in an increase in the total number of Tj^+^ cells (Fig. S1H’, and M) although to a lesser extent than *Tj-Gal4*, suggesting that signals from the hub may also partially influence CySC proliferation. Given the stronger phenotypic effect observed with *Tj-Gal4*, we proceeded with this driver line for subsequent experiments.

**Figure 2.**
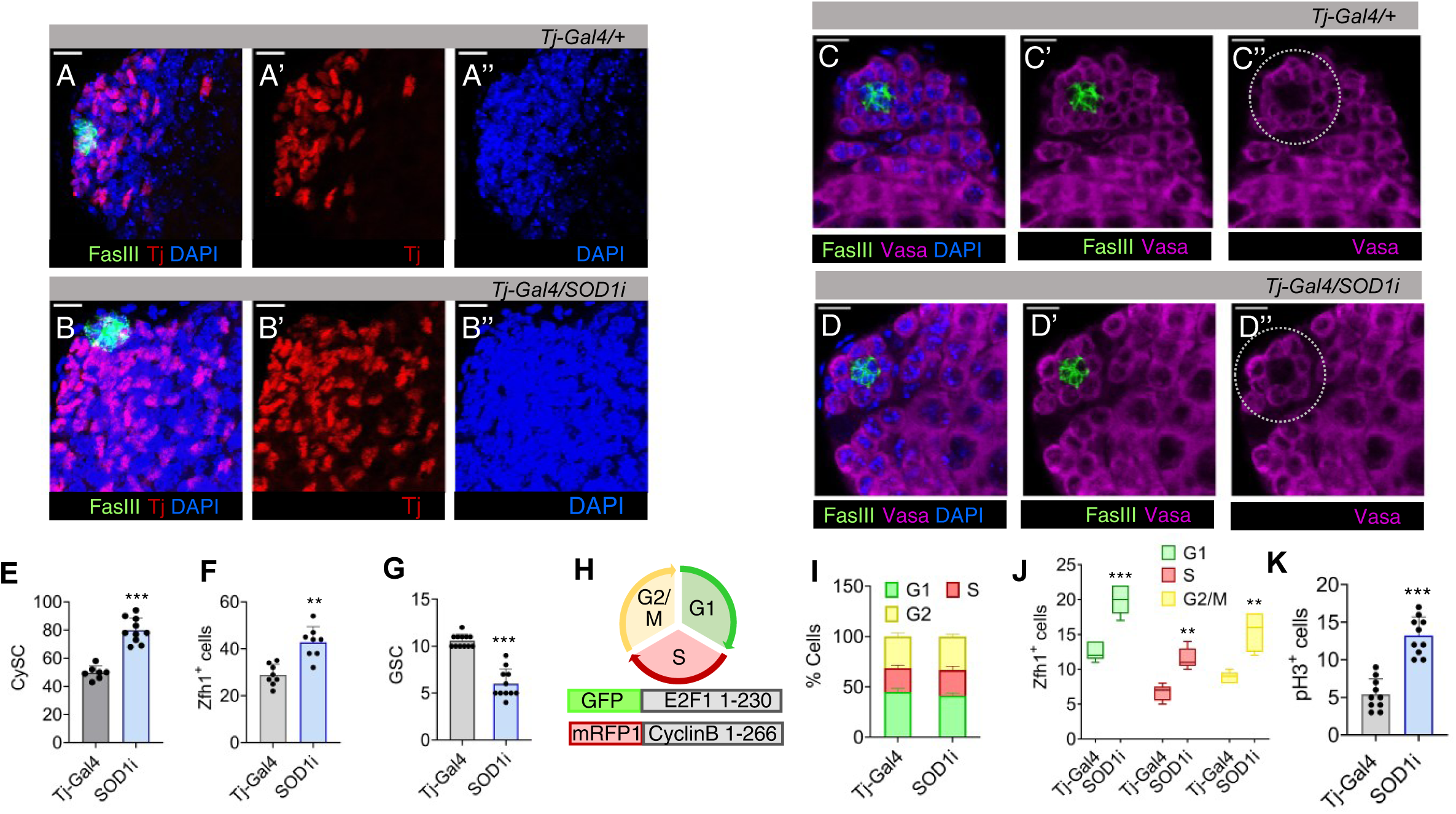
Redox disequilibrium in Cyst stem cell lineage deregulates early CySCs and decreases the GSC number. (A-B) Distribution of Tj^+^ CySC/early cyst cell around the hub (FasIII) in control and Tj driven *Sod1RNAi* testis. (C-D) Represents the rosette arrangement of Vasa^+^ GSCs flanking the hub (dotted area). (E-G) Mean number of Tj^+^ cells, Zfh1^+^ early CySCs and GSCs in control and *Sod1RNAi* lines shown as mean ± s.e.m, n = 10, *** *P(unpaired t-test)<0.0001, **P(unpaired t-test)<0.0001.* (H) Construct design of the FUCCI reporter line for tracking the cell cycle stages in *Drosophila* tissues, containing degrons of Cyclin B and E2F1 proteins fused with RFP or GFP. G1, S and G2/M is represented by GFP^+^, RFP^+^ and dual labelled cells respectively. (I-J) Comparative changes in the cell cycle phases (I) or Zfh1^+^ cells (J) present in G1, S and G2/M phase at the niche zone between control and *Tj>Sod1i*. Data denote mean ± s.e.m, n = 10, *** *P(unpaired t-test)<0.0001, **P(unpaired t-test)<0.001.* (K) Mean number of pH3+ cells, a mitotic marker. Data points denote mean ± s.e.m, n = 10. Scale bar - 10 µm. See also Figure S2.

The enhancement in the number of CySCs was also reflected by ∼2-fold change in Zfh1^+^ cells (Fig. 2F, S2R(I)-S(I)). However, Vasa^+^ cells flanking the hub (GSCs) showed a substantial reduction in number (Fig. 2C’’-D’’ and G). This decline was further validated by western blot analysis, which revealed lower overall detectable levels of Vasa protein, indicating a potential impact of the altered redox state on germline maintenance (Fig. S2N). Since, Tj shows a minor expression in hub cells, we validated our phenotypic data using *C-587-Gal4* that is specific for CySCs and CCs. Similar to *Tj>Sod1*, *C-587-Gal4>Sod1i* also showed a reduction in the number of neighbouring GSCs (Fig.S1N). The observations were further verified using *Sod1i* localized in different chromosome to avoid any chromosome or balancer-based biasness and a similar result was obtained where alterations in CySC ROS affected GSC number (Fig. S1I-L and O-P). However, the phenotypic effect of higher ROS was limited to its origin in CySCs only because ablation of GSC redox status did not cause any marked change in niche composition. *Nos-Gal4* driven knock-down of Sod1 in GSCs did not result in significant change in CySC number (Fig. S2A’, B’ and E) but effected a considerable reduction on Vasa^+^ cells (Fig. S2C’, D’ and F), aligning with previous observations^17^. Although Sod1 is the predominant enzyme which also localizes in the intermembrane space of mitochondria, we observed a similar result upon depleting the matrix localized Sod2 in both GSCs and CySCs (Fig. S2G-M), indicating the involvement of mitochondrial ROS for the observed cellular phenotypes.

To confirm the active proliferation of cells and rule out the possibility of arrested growth, we utilized fly-Fluorescent Ubiquitination-based Cell Cycle Indicator (FUCCI) system^41^ which comprises of two reporter constructs marking G1/S transition (green), S (red) and G2/early mitosis (yellow) (Fig. 2H). Testes from *Tj>Sod1i* exhibited an approximately two-fold increase in cell number across all stages of the cell cycle (Fig. S2O). However, the percentage of cells in each phase remained unchanged (Fig 2I), indicating that the observed increase in different phases is a consequence of more cells entering proliferation rather than alterations in the duration of different cell-cycle stages. The enhanced number of dividing cells was contributed mainly by progressive division of CySCs, showing >2-fold difference in the accumulation of Zfh1^+^ nuclei in S and G2/M phases (Fig. 2J, S2RV-VII and SV-VII). The number of Zfh1 labelled cells in G1/S transition was also substantially more (Fig. S2RIII and SIII), indicating dysregulated redox ensued an increased mitotic index of CySCs. The enhanced mitotic activity of these proliferating niche cells is further supported by increased phospho-histone H3 (PH3) incorporation, indicating a higher mitotic frequency compared to the control (Fig. 2K, S2T-U), along with elevated cyclin D expression (Fig. S2Q). Downregulation of Sod1 in GSCs through *Nos-Gal4* demonstrated no significant change in G1/S, S and G2/M phase numbers, confirming our previous observation that altering ROS levels in GSCs does not have any non-autonomous effect on CySCs numbers (Fig. S2P).

### CySC induced ROS couples precocious differentiation of stem cell lineages

Since in *Tj>Sod1i* testes, the increased number of Tj*^+^* cells, marking CySCs and early differentiating cells, exceeded the number of CySC-specific Zfh1*^+^* cells (Fig. 2J and S2O), we checked for parallel enhancement of cellular differentiation using the corresponding marker, *Eya* (Eyes absent).

We found that Eya^+^ cells clustered towards the hub, along with a significant increase in their number in Sod1 depleted CySCs (Fig. 3A, B and D). This increased number of Eya^+^ cells does not suggest that these differentiating cells are proliferative. Instead, we propose that the knockdown of Sod1 may alter the timing or regulation of cyst cell differentiation, leading to an accumulation of Eya^+^ cells near the niche. The supposed premature expression of Eya resulted in a population of differentiating cyst precursor cells co-expressing Eya-Zfh1 (Fig. 3A’’, B’’ and D). This population was found to be deviating from the normal partitioning of wild-type representative populations (Fig. 3C). In contrast, depletion of *Sod1i* in the germline lineage using *Nos-Gal4* did not result in a similar phenotypic arrangement (Fig. S3A-D). These findings suggest that alterations in the redox state of GSCs can impact their numbers, even when the changes are induced non-autonomously through CySCs^17^.

**Figure 3.**
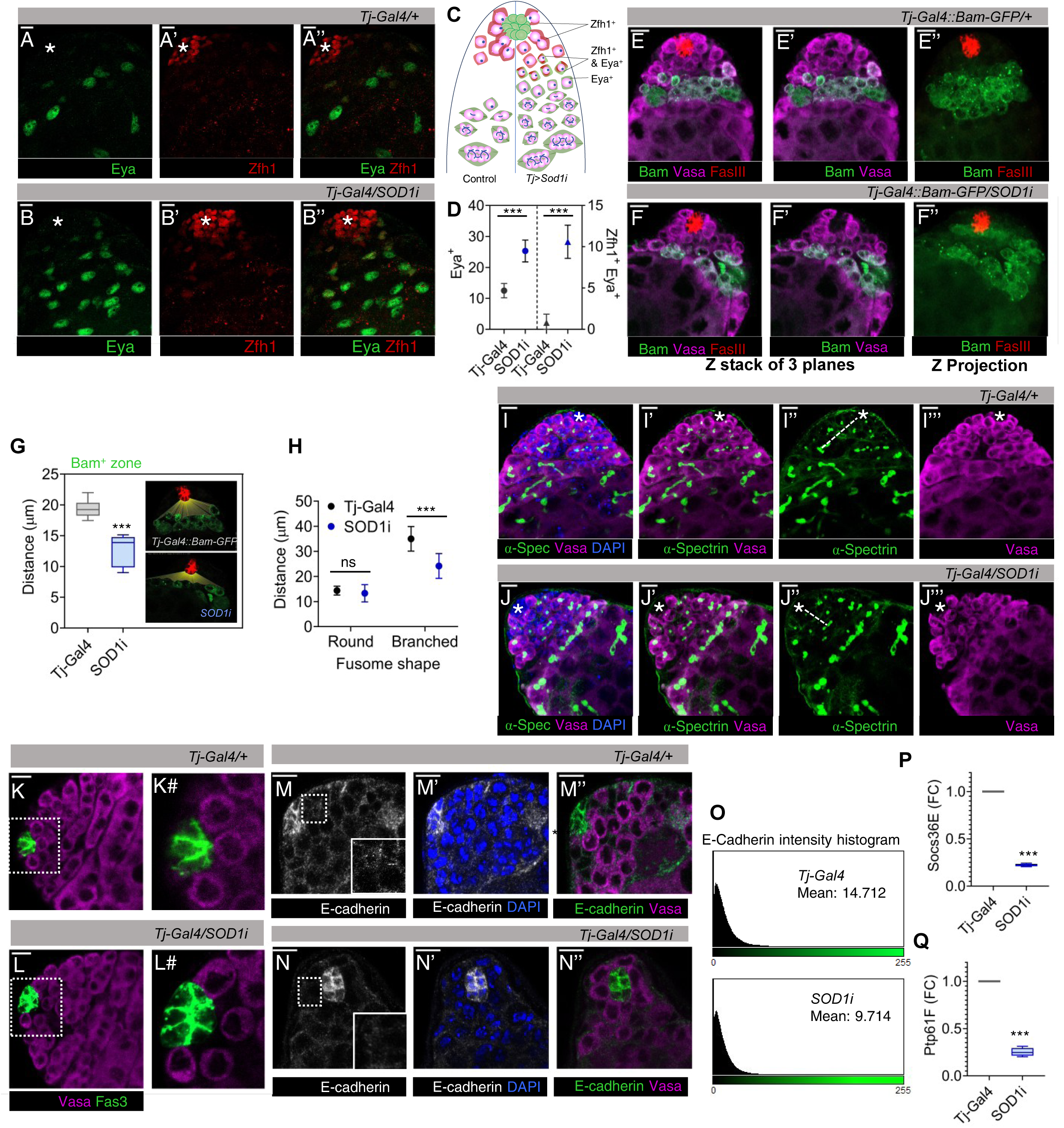
ROS imbalance in CySCs promotes differentiation of both GSCs and CySCs. (A-D) Number of early CySCs (Zfh1^+^) (A’-B’), late differentiating CCs (Eya^+^) (A - B) and Zfh1, Eya co-expressing population (A’’-B’’) were imaged, represented schematically (C) and quantified with respect to control and *Sod1RNAi* lines (D). Data points represent mean ± s.e.m, n = 10, *\*\*\*P(unpaired t-test)<0.0001*. (E-F) The effect of Sod1 depletion in CySCs on the differentiation status of GSCs as observed using *Bam-GFP* reporter line. (G) The relative distance of the differentiation initiation zone (Bam^+^) from the hub (red) in control and *Sod1i* testis quantified in single sections and shown as mean ± s.e.m, n = 10, *\*\*\*P(unpaired t-test)<0.0001.* (H-J) The shape and size of the spectrosomes marked with α-spectrin (green) were imaged (I’’-J’’) and quantified for their distance from the hub (marked with asterisk) (H), n = 10, *\*\*\*P(unpaired t-test)<0.0001*. Branched fusome marks differentiating populations (I’-J’). (K-L) Represents the rosette arrangement of Vasa^+^ GSCs flanking the hub. The digitally magnified region around the hub (dotted line) as seen in control (K#) and *Sod1RNAi* (L#). (M-N) Comparative staining of CySC-GSC contacts through adherens junction using E-cadherin (monochrome/green). Dotted area near the hub has been expanded as inset to show loss of E-cadherin network. (O) E-cadherin intensity histogram plot generated from ImageJ representing the mean intensity of expression in the region flanking the hub and GSCs. (P-Q) Fold change in expression of Stat dependent transcripts Socs36E (P) and Ptp61F (Q) among control and *Sod1i* niche, obtained through qPCR. Data is shown as mean ± s.e.m, n = 3, *\*\*\*P(unpaired t-test)<0.0001*. ns – not significant. Scale bar - 10 µm. See also Figure S3.

Together with reduction in Vasa^+^ GSCs shown earlier (Fig. 2G), we also detected gonialblast expressing the differentiation-promoting factor Bam, to be much closer to hub in *Tj>Sod1i* testes (Fig. 3E’’ and F’’). The mean distance between the hub and Bam^+^ cells was found to be shortened by ∼7 microns (Fig 3G). The relative advancement of spermatogonial differentiation towards the hub was also ascertained by tracking the transformation of GSC-specific spectrosome into branched fusomes connecting the multi-cell gonialblast (Fig. 1A). The branching fusomes were found more proximal to hub than controls (Fig. 3H, I’’ and J’’), along with parallel reduction in number of spectrosomes, implying a decline in early-stage germ cells (Fig. S3F). This loss could possibly be attributed to loss of intercellular contacts, particularly with hub cells (Fig. 3K# and L#). The observation corroborated with decreased expression pattern of E-cadherin flanking the hub and GSCs, in *Tj>Sod1i* (Fig. 3M-O) and could be one of the reasons for GSCs dissociation and differentiation (Fig. 3L#), Disengagement of GSCs from the hub caused reduction in Socs36E and Ptp61F transcripts which subsequently alter Stat expression, a key driver of E-cadherin expression (Fig. 3P and Q). These results suggest that disruption of intercellular redox gradient affected the premature differentiation of stem cells in the niche.

### Disrupted niche architecture compromises GSC-CySC communication

GSCs and CySCs intercommunicate through several factors such as EGFR, PI3K/Tor, Notch^42^ to support CySC maintenance^43^. EGF ligands secreted by spermatogonia maintain differences in cell polarity and segregates self-renewing from differentiating populations^44^. *Tj>Sod1i* cells showed lower levels of cell-polarity marker Dlg (Fig. 4A”, B” and E), that expanded more into the differentiated zone (Fig. S4A), indicating loss of cell polarity and aligning with the overtly proliferative nature of these cells. Along with Dlg, a net reduction in pErk levels was observed across sections in *Sod1i* testis (Fig. 4C’, D’ and F). This reduction in overall Erk levels may be attributed to the loss of cell contact and altered GSC-CySC balance in *Sod1i* testis. The proliferation of cyst cells correlated with increased levels of self-renewal inducers, including the elevated expression of Hedgehog (Hh) pathway components^32^, in *Tj>Sod1i* testis. The depletion of Sod1 in CySC resulted in elevated expression of Hh receptor, Patched (Ptc) (Fig. 4I, S4V) which extended to regions farther from the hub, overlapping with expanded Tj^+^ cells (Fig. 4G and H). Since, Ptc itself is a transcriptional Hh target, its expression in CySCs suggests active Hh signalling^45^, further supported by higher levels and a similar pattern of Hh transcriptional effector, Cubitus interruptus (Ci), in *Sod1i* testis (Fig. 4M’’, N’’, J and S4W). Since, Ptc, Ci, and a GPCR-like signal transducer Smoothened (Smo) are all transcriptionally driven by Hh, and Hh itself is regulated by higher levels of Ci^46,47^, we expectedly found higher levels of transcripts for these pathway genes (Fig. S4C-F). The overt proliferation of CySCs under high ROS conditions was suppressed by reducing hh mRNA levels in hub cells and CySCs using *Hh-RNAi* (Fig. 4L and Fig. S4G-I), which also rescued the GSC population (Fig. 4K and S4J-L). The levels of Sod1 transcripts under depletion of Sod1 alone and Sod1, hh double knock-down were almost similar (Fig. S4M). To determine whether the changes observed in the germ cell population were a sole consequence of *Sod1* knockdown in CySC, we manipulated CySC numbers using *UAS-Ci* and assessed their impact on neighbouring GSCs. Ci overexpression driven by *Tj-Gal4* led to a higher number of Tj^+^ cells (Fig. S4N, R’’ and S-U) and a complete loss of Vasa^+^ GSCs (Fig. S4O, and R’’’). However, the effect on CySC and GSC population was less severe when driven by *C-587 Gal4* (Fig.S4N, O, Q’’and Q’’’).

**Figure 4.**
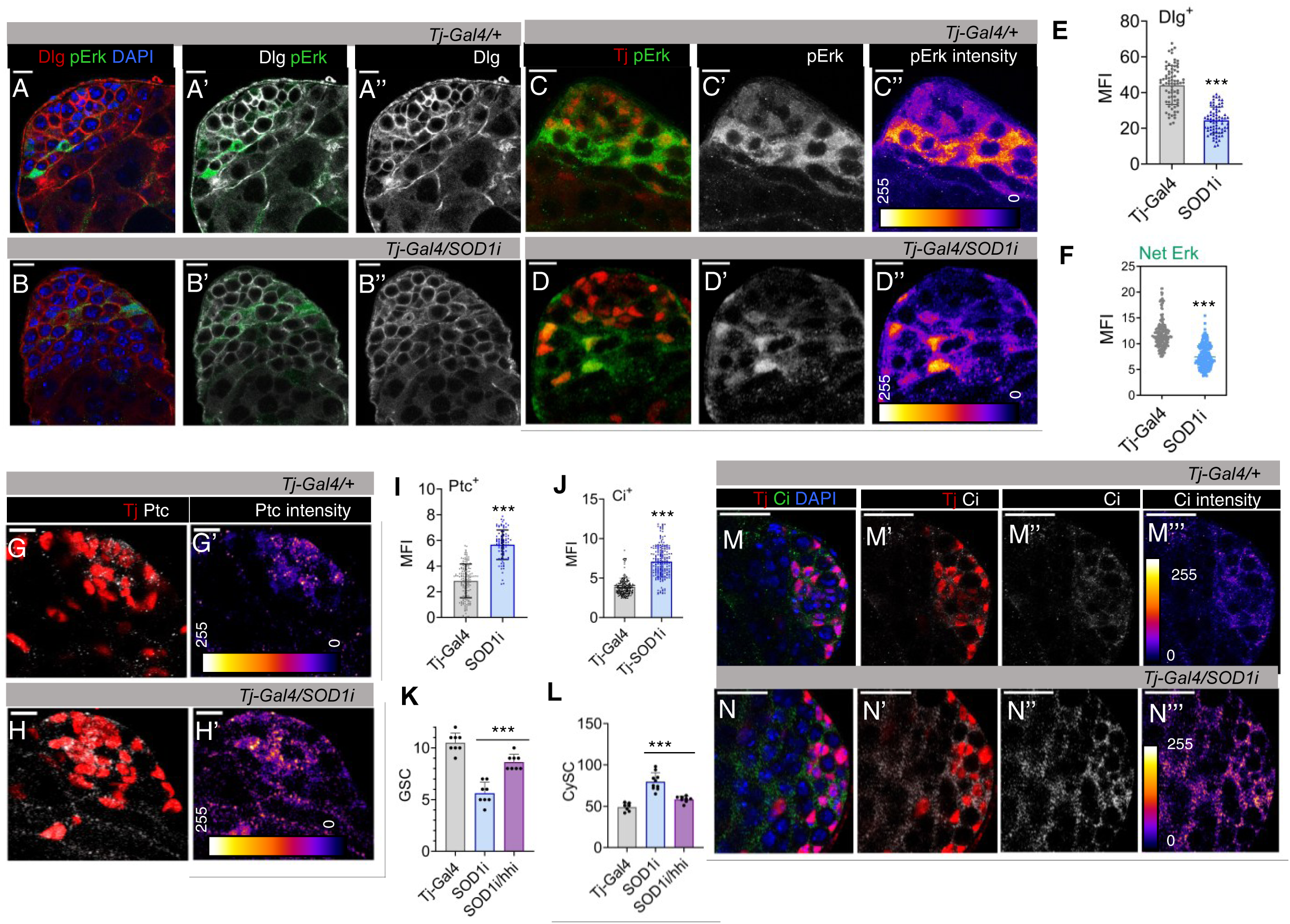
Altered CySC redox state affects niche maintenance signals. (A-B) The expression of cell polarity marker Dlg (red) in control and *Sod1i* stem-cell niche (A’’-B’’), (C-D) Representative image showing pErk distribution parallelly with Tj^+^ cells and its expression pattern through Fire LUT (C’’-D’’). Scale bar - 10 µm. Mean florescence intensity (MFI) corresponding to Dlg level (E) was quantified, n = 75, *\*\*\*P(unpaired t-test)<0.0001.* (F) MFI of total pErk expression was quantified, n =200, *\*\*\*P(unpaired t-test)<0.0001*. (G’-H’) Representative image showing distribution of Patched (Ptc) in Tj^+^ cells in control and *Tj>Sod1i*. (I-J) Quantification of Ptc (I) and Hh effector Ci (J) expression through fluorescence intensity as mean ± s.e.m, n (Ptc) = 90, n (Ci) = 200, *\*\*\*P(unpaired t-test)<0.0001* (K-L) The number of Vasa^+^ (K) and Tj^+^(L) cells across control, *Sod1i*, and *Sod1i/Hhi* rescue samples depicted as mean ± s.e.m, n = 10, *\*\*\*P(unpaired t-test)<0.0001*. See also Figure S4.

### Elevating CySC antioxidant defence promotes GSC self-renewal

To further substantiate the role of ROS in coordinating the stem cell populations, we strengthened the cellular defences against oxidants by overexpressing Sod1 in CySCs. We checked the overall ROS level using DHE (Fig. S5I) and monitored the relative GSC/CySC populations. Together with reduction in superoxide levels, we observed an increase in the number of Vasa^+^ cells (Fig. 5A and S5A’’-B’’), and a slight reduction in Tj^+^ cells (Fig. 5B and S5E’’-F’’). Any morphological anomalies associated with cell-cell adhesion or abnormal cellular dispersion, was not observed (Fig. S5A-B and E-F). Enhancing the levels of Sod1 in GSCs promoted the growth of GSC-like cells (Fig. S5C’’, D’’ and J) but did not have any prominent effect on CySCs (Fig. S5G’’, H’’ and K). The increment in Vasa^+^ GSCs under CySC-induced low redox conditions was also represented by a parallel increase in spectrosome number (Fig. 5C-E). The uptick in GSC number due to scavenging of ROS in CySC, might be a result of delayed differentiation, as indicated by displacement of Bam^+^ zone away from hub; thus, confirming non-cell-autonomous role of CySC ROS in maintaining GSC fate (Fig. 5F-H). The data suggest that reducing redox profile of CySCs differentially affected their own self-renewing propensities along with GSC maintenance and optimum redox state of both the stem-cell populations is controlled by CySCs.

**Figure 5.**
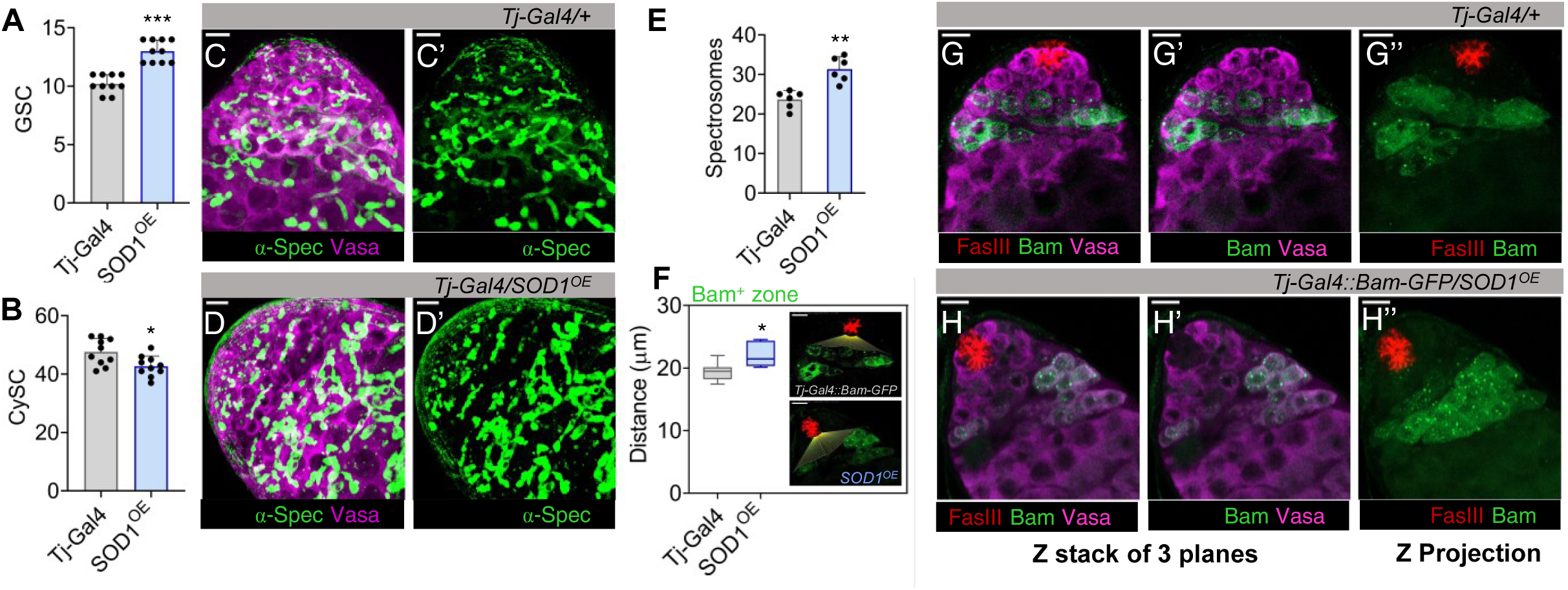
Low CySC ROS sustains GSC maintenance. (A-B) Bar graphs illustrating variations in Vasa^+^ GSC (A) and Tj^+^ CySCs (B) numbers upon overexpressing Sod1 (*Tj>Sod^OE^*), represented as mean ± s.e.m, n = 10, *\*\*\*P(unpaired t-test)<0.0001, *P(unpaired t-test)<0.01*. (C-E) The spectrosomes labelled with α-spectrin (α-spec) were imaged (C’-D’) and their number was quantified (E), n = 10, *\*\*P(unpaired t-test)<0.001*. (F-H) The initiation of gonialblast differentiation was determined by measuring the distance of Bam^+^ zone from hub (red), n = 10, *\*P(unpaired t-test)<0.01* (F), (G’’-H’’) shows distribution of Bam^+^ reporter expressing cells. Scale bar - 10 µm. See also Figure S5.

**Figure 6.**
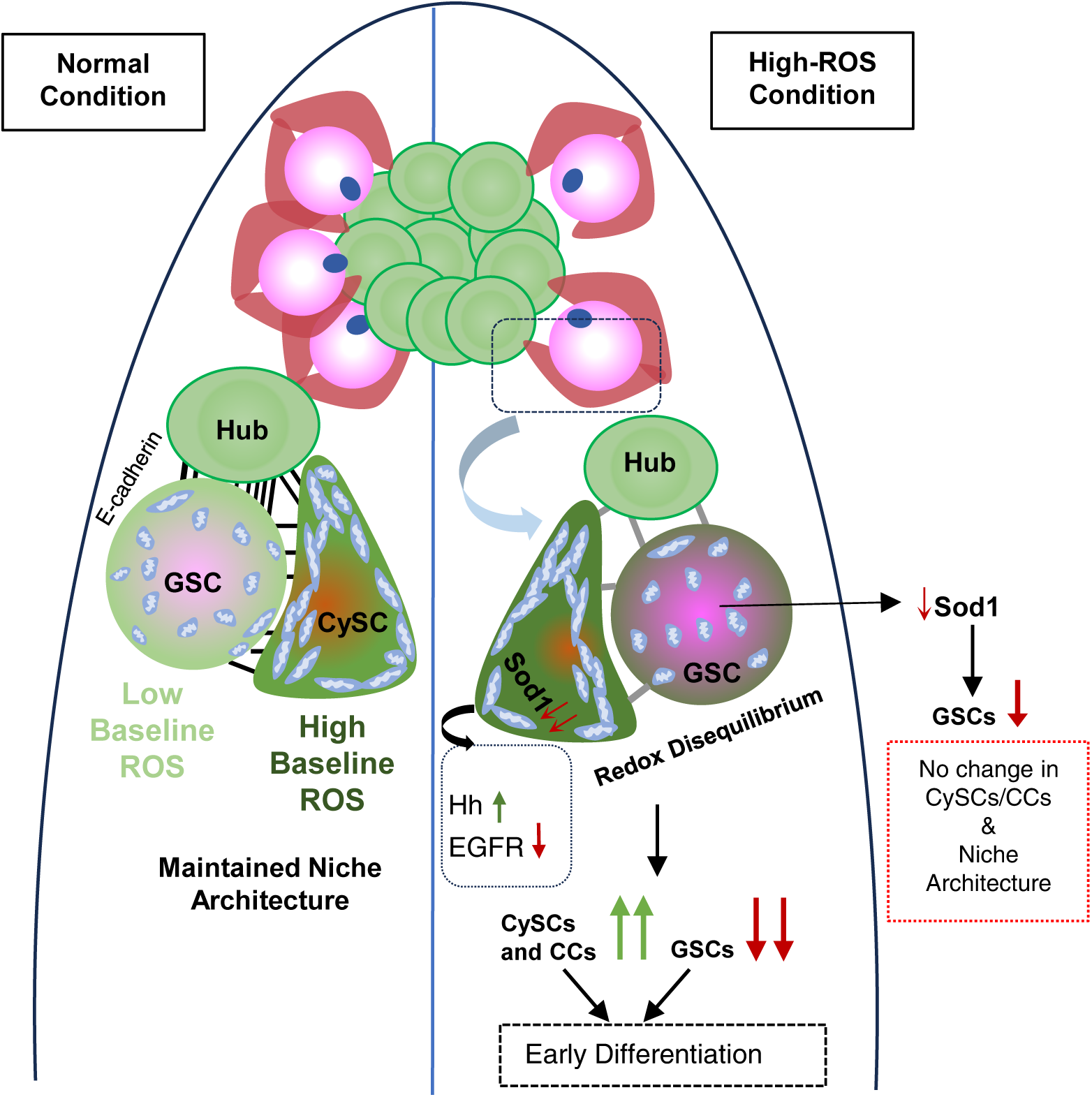
Proposed role of intercellular redox balance in germline maintenance and differentiation. GSC and CySC are associated with different mitochondria abundance which correspond to the presence of a redox differential between two stem cell populations. Disequilibrated redox potential among the two populations due to depletion of Superoxide dismutase, enhances CySC proliferation and precocious GSC differentiation, thereby disrupting the homeostatic balance between the two stem populations.

## Discussion

ROS is a crucial mediator of stem cell maintenance where these species act as second messengers to induce different post-translational protein modifications thereby, affecting cell fate^48,49^. The effect of ROS is mainly considered to be autocrine in nature, where generation of these oxidative species is usually restricted to specific cellular compartments, ensuring they act close to their targets. Recent studies have reported non-autonomous generation of ROS by NOX enzymes or upon stimulation by growth factors, to play a critical role in regenerative growth^50–52^. However, in either case, the target cell serves as both the source and the site of response. Among ROS, hydrogen peroxide (H₂O₂) stands out due to its stability and membrane permeability^53^, allowing it to diffuse from its source^54^. Non-myocytic pericardial cells (PCs) in *Drosophila* exhibit elevated ROS that act in a paracrine manner to influence adjacent cardiomyocytes (CMs) not by direct diffusion but through activation of the D-p38 MAPK cascade within PCs^55^. In cardiomyocyte monolayers, wounding induces H_2_O_2_ and superoxide accumulation that propagate via gap junctions, creating a cell-to-cell ROS wave where even distant cells show increased cytosolic H_2_O_2_ and altered proteomes^56^. Together, these findings suggest that ROS can spread between cells either by direct diffusion/gap-junctional transfer or indirectly by modulating signaling cascades and secreted factors. Our findings indicate that CySCs due to their high baseline ROS levels, affect the redox state of their neighbourhood including GSCs. ROS imbalance in CySCs enhance the oxidative levels of their surrounding that is, GSC that cause them to differentiate. High ROS mediated differentiation of GSCs has already been reported earlier^17^ and depletion of superoxide dismutases in GSCs did result in reduction of their number. However, perturbation of ROS in GSCs did not have significant effect on CySC population. This shows that cell non-autonomous effect of ROS is observed only when it originates from CySCs.

Our observations in *Drosophila* testicular stem cell niche suggested a previously unaccounted role of superoxide dismutases in maintaining niche homoeostasis. The redox perturbations affected cyst stem cell dynamics which indirectly influenced the germline. The higher ROS threshold of CySCs maintained the physiological redox state of GSCs. Reduced ROS in GSC was accompanied with its proliferation^57^, phenocopied when Sod1 was overexpressed in its neighbours (Fig. 5). However, we feel that net alterations in GSC state were the combinatorial effect of oxidative stress and niche alterations brought about by CySC deregulation.

GSCs are anchored to the hub cells via adherens junctions, allowing them to receive crucial maintenance signals^28^. These include Bone Morphogenetic Proteins (BMPs) such as Decapentaplegic (Dpp) and Glass bottom boat (Gbb), which activate the BMP pathway^31^ required for GSC self-renewal, and Unpaired (Upd), which triggers the JAK-STAT pathway to promote GSC adhesion to the hub^58^. The suppression of these pathways activates Bam, acting as a switch from transit-amplifying cells to spermatocyte differentiation^59,60^. Elevated ROS in CySC possibly affected STAT phosphorylation through S-glutathionylation^61,62^, reducing the levels of STAT-dependent transcripts such as Socs36E^63^, Ptp61F and E-cadherin. This depletion compromises GSC adhesion, resulting in detachment from neighboring cells, similar to E-cadherin loss conditions ^58,64–66^. This resulted in compromised GSC-CySC paracrine receptions such as EGFR which play a crucial role in the maintenance of cell-polarity that segregates self-renewing CySC populations from the ones receiving differentiation cues ^44,67,68^, leading to the accumulation of cells co-expressing both stemness and differentiation markers. In contrast to GSCs, ROS imbalance in CySCs induced accelerated proliferation as well as differentiation due to deregulation of EGFR and Hedgehog pathways. Although, redox modulation of EGFR pathway components has been previously demonstrated^69^, in this study we found Hedgehog to be probably susceptible to redox regulation. The redox dependent induction of Hedgehog probably, contributed to enhancement in CySC numbers which partly contributed to depletion of GSCs.

However, we do not rule out the possibility of hub cells playing an equivalent role in GSC maintenance, given their parallel effect in suppressing GSC differentiation^30^. Tj majorly expresses in CySCs but also shows modest expression in hub. Therefore, the possible depletion of Sod1 in both hub and CySCs by *Tj-Gal4* together with, demonstration of a stronger phenotype in *Tj-Sod1RNAi* testis compared to CySC specific *C587-Sod1RNAi*, indicated potential hub contributions in CySC self-renewal. While ROS accumulation typically enhances EGFR signaling promoting premature GSC differentiation^70^, the loss of EGFR in somatic cells increases the number of GSCs^67^ and is associated with fewer somatic cells ^71^. However, we observed that Sod1 depletion caused parallel reduction of both pErk levels and GSC number. It is quite possible that the response of germline to Erk modulation is more context-dependent which is influenced by other receptor tyrosin kinases beyond EGFR. To test this hypothesis, we had tried an EGFR gain-of-function rescue of CySC depletion but the driven progenies were lethal in the absence of Sod1. We also avoided usage of Gal80 dependent clonal populations to maintain a homogenous genetic background and prevent false readouts due to diffusion of ROS signals in unaltered neighbourhood.

The proposed concept of intercellular ROS communication can be of importance in deciphering the biochemical adaptability and plasticity of different niches influencing stem cell fate, ensuring niche size and architecture to prevent stem cell loss and aging. This has been observed in neural stem cells where inflammation-induced quiescence recovers the regenerating capacity of aging brain^72^. In addition to metabolic variations, dependence of the germline on somatic neighbours for its redox state might be one the reasons behind its presumed immortal nature and resistance to aging^73^. The same can also be applied to cancer stem cells, which maintain niche occupancy and resist oxidative stress-induced apoptosis, probably by receiving redox cues from the environment. However, further work is required to elucidate the myriads of driver mechanisms intersecting in the realm of redox regulation that extend beyond the present system into broader translational areas.

## Materials and Methods

### Fly strains

Fly stocks were maintained and crosses were set at 25°C on normal corn meal and yeast medium unless otherwise indicated. All fly stocks (BL-24493) *UAS SOD1 RNAi*, (BL-32909) *UAS SOD1 RNAi*, (BL-29389) *UAS SOD1 RNAi*, (BL-32983) *UAS SOD2 RNAi*, (BL-24754) *UAS SOD1*, (BL-32489) *UAS hhRNAi*, (BL-25751) *UAS Dcr-nos Gal4*, (BL-55122) *UAS-FUCCI*, were obtained from Bloomington Drosophila Stock Centre. *Nos-Gal4*, *Tj-Gal4*, *Tj-Gal4-BamGFP*, *gstD1-GFP*, *TFAM-GFP, UAS-Ci* were kind gift from U. Nongthomba, P. Majumder, K. Ray, B. C. Mandal, Hong Xu, LS Shahidhara labs respectively. Parents were maintained at 25°C, and crosses were set at the same temperature. To ensure optimal Gal4 activity, progenies were shifted to 29°C until eclosion, with aging also conducted at 29°C. Males aged 3–5 days were used for all experiments. The control lines for all experiments are the corresponding Gal4-line crossed with wild type *OregonR^+^*.

### Dissection and immunostaining

Anesthetized flies were dissected in 1X Phosphate Buffer Saline (PBS). All incubations were carried out at room temperature (25°C) unless otherwise mentioned. Testes were fixed in 4% paraformaldehyde for 30 minutes, followed by three washes with 0.3% PBTX (PBS + TritonX 100). Post-blocking in 0.5% Bovine Serum Albumin, testis was incubated overnight at 4°C for primary antisera, followed by washing in 0.3% PBTX 3 times 15 minutes each before incubating with secondary antibody. The tissues were counterstained with DAPI (4,6-diamidino-2-phenylindole) for 20 minutes, followed by three washes in 0.3% PBTX for 15 minutes each. For Patched antibody staining, a modified protocol utilizing PIPES-EGTA buffer was used as described^74^. For Eya staining, the tissues were incubated in primary antibody for 2 days. The dpErk was labelled by dissecting fly testes in Schneider’s media, followed by fixation in 4% PFA for 30 min, 3x wash in 0.1% PBTX, blocked and incubated in primary antibody overnight; each step supplemented with phosphatase inhibitor cocktail 2 (1:100, Sigma, cat#P5726). Samples were mounted in DABCO anti-fade medium prior imaging.

The following primary antibodies were used in the experiments - anti-FasIII (7G10) (1:120, DSHB (Developmental Studies Hybridoma Bank)), anti-Eya (1:50, DSHB), anti-DEcad (DCAD2) (1:50, DSHB), anti-α-Spectrin (3A9) (1:20, DSHB), anti-Ptc (1:50,DSHB), anti-Ci (1:50, DSHB), anti -Dlg (1:50), anti-βtubulin (1:300, DSHB), anti-pERK (1:100, Cell Signaling (4370)), ATP5A (1:700, Abcam (ab14748)), anti-Tj (1:5000), anti-Vasa (1:4000), anti-Zfh1(1:2000). The primary labelling was detected using appropriate Alexa-Flour tagged secondary antibody (ThermoFisher Scientific).

To assess superoxide levels, testes were dissected in Schneider’s medium (SM) and immediately incubated in 30 µM Dihydroxyethidium (DHE) at 25 °C in the dark. Testes were washed thrice with SM for 7 minutes each before quantifying the emitted fluorescence using SpectraMax iD5 (Molecular Devices).

### Quantitative Reverse transcription–PCR

For semi-quantitative RT-PCR and qRT-PCR analyses, total RNA was isolated from testes of 3-5 days old male flies using Qiagen RNA extraction kit. RNA pellets were resuspended in nuclease-free water and quantity of RNA was spectrophotometrically estimated. First-strand cDNA was synthesized from 1-2 µg of total RNA as described earlier. The prepared cDNAs were subjected to real time PCR using forward and reverse primer pairs as listed below, using 5 µl qPCR Master Mix (SYBR Green, ThermoFisher Scientific), 2 pmol/µl of each primer per reaction for 10 µl final volume in ABI 7500 Real time PCR machine. The fold change in expression was calculated through 2^-ΔΔCt^ method. The primers used are listed in Supplementary Table S1.

### Imaging and image analysis

Confocal imaging was carried out using Zeiss LSM 900 confocal microscope, using appropriate dichroics and filters. Images were further analyzed and processed for brightness and contrast adjustments using ImageJ (Fiji). Mean intensity measurements were carried out using standard Fiji plug-ins,The relative distance from the two reference points in the images was estimated through Inter-edge Distance ImageJ Macro v2.0 from Github. All images were assembled using Adobe Photoshop version 21.2.1.

### Quantification of GSCs, CySCs and CCs

For GSC quantification, only cells in direct contact with the hub were considered. While single slices are shown for representation, counting was performed on individual slices of z-stacks using the semi-automated Cell Counter plugin in Fiji for all specified genotypes.

For CySC quantification, all Tj-positive and Zfh1-positive cells present in each testis were counted and represented. For cyst cells quantification, only those near the hub were considered, which does not reflect the total number of Eya-positive cells. All quantifications were performed using the Cell Counter plugin in Fiji.

#### Mitochondrial and *gstD1* -GFP quantification

The mitochondrial analyzer first generates 2D mitochondrial regions of interest (ROIs), followed by the measurement of their total area and circularity. Initially, this is done using a thresholded image of a single slice. For *gstD1-*GFP quantification, we have quantified the region of *gstD1* - GFP that overlapped with Vasa-positive germ cells which are in direct contact with hub to assess the presence of ROS reporter signal within the germline compartment. To ensure accurate cell boundary demarcation, we used Dlg staining as an additional parameter.

### Immunoblot Analysis

Protein was extracted from the dissected testes using RIPA buffer as described previously and quantified using Bradford reagent (Biorad). Equivalent concentration of lysate was denatured using 1X Laemelli Buffer with 1 M DTT at 95°C, separated in 10% SDS-PAGE and transferred onto PVDF membranes. Membranes were blocked with commercial blocking solution in TBST base before sequential primary antibody incubation with anti-Vasa and anti-β-Tubulin (DSHB, E7). Secondary detection was performed using HRP-tag anti-mouse antibody (GE Amersham).

### Statistical analyses

The sample sizes carrying adequate statistical power are mentioned in the figure legends. Statistical significance for each experiment was calculated using two-tailed Student’s t-test, unless otherwise mentioned, using GraphPad Prism 8 software. The *P* values were calculated through pairwise comparison of the data with wild type or driver alone and driven RNAi lines. Significance values for sample sizes mentioned in figure legends were represented as * *P* < 0.01, ** *P* <0.001, or *** *P* < 0.0001.

## Author contribution

D.S. and O.M conceived the study and designed the experiments. O.M, A.C and T.C performed the experiments. D.S and O.M analyzed the data. D.S, O.M. and A.C. wrote the manuscript.

## Acknowledgements

We thank P. Majumder, U. Nongthomba, K. Ray, G. Ratnaparkhi, B. C. Mandal, Hong Xu lab and S. C. Lakhotia for kindly sharing some of the transgenic fly lines. Christian Bökel for sharing experimental protocols. Anti-Tj antibody was a generous gift from Prof. D. Godt, University of Toronto. Anti-Vasa and Anti-Zfh1 were kindly gifted by Prof. R. Lehmann, New York University. We also thank Dr. Bama Charan Mandal for generously sharing his Confocal microscope imaging system and Sudeshna Majumder for preliminary data. We acknowledge BDSC for fly stocks and DSHB for antibodies. This work is supported by Innovative Young Biotechnologist Award by Department of Biotechnology (BT/12/IYBA/2019/01), Early Career Research Award by Science and Engineering Research Board (ECR/2018/000009), BHU-Institute of Eminence grant (R/Dev/IoE/Incentive/2021-22/32452). The authors also acknowledge financial support from DST-INSPIRE fellowship (to O.M.) and UGC-CAS doctoral fellowship (to T.C.).

## Declaration of interest

The authors declare no competing interests.

## Data availability

The data that support the findings of this study are available within the main text and its Supplementary Information file. The lead contact will share all raw data associated with this paper upon request. This study did not utilize or generate any unique datasets or codes.

## Notes

### Competing Interest Statement

The authors have declared no competing interest.

### Summary of Updates

The manuscript has been revised after addressing reviewers comments.

