## Supplementary Figures for "Superoxide Dismutases maintain niche homeostasis in stem cell populations"

**Figure S1**

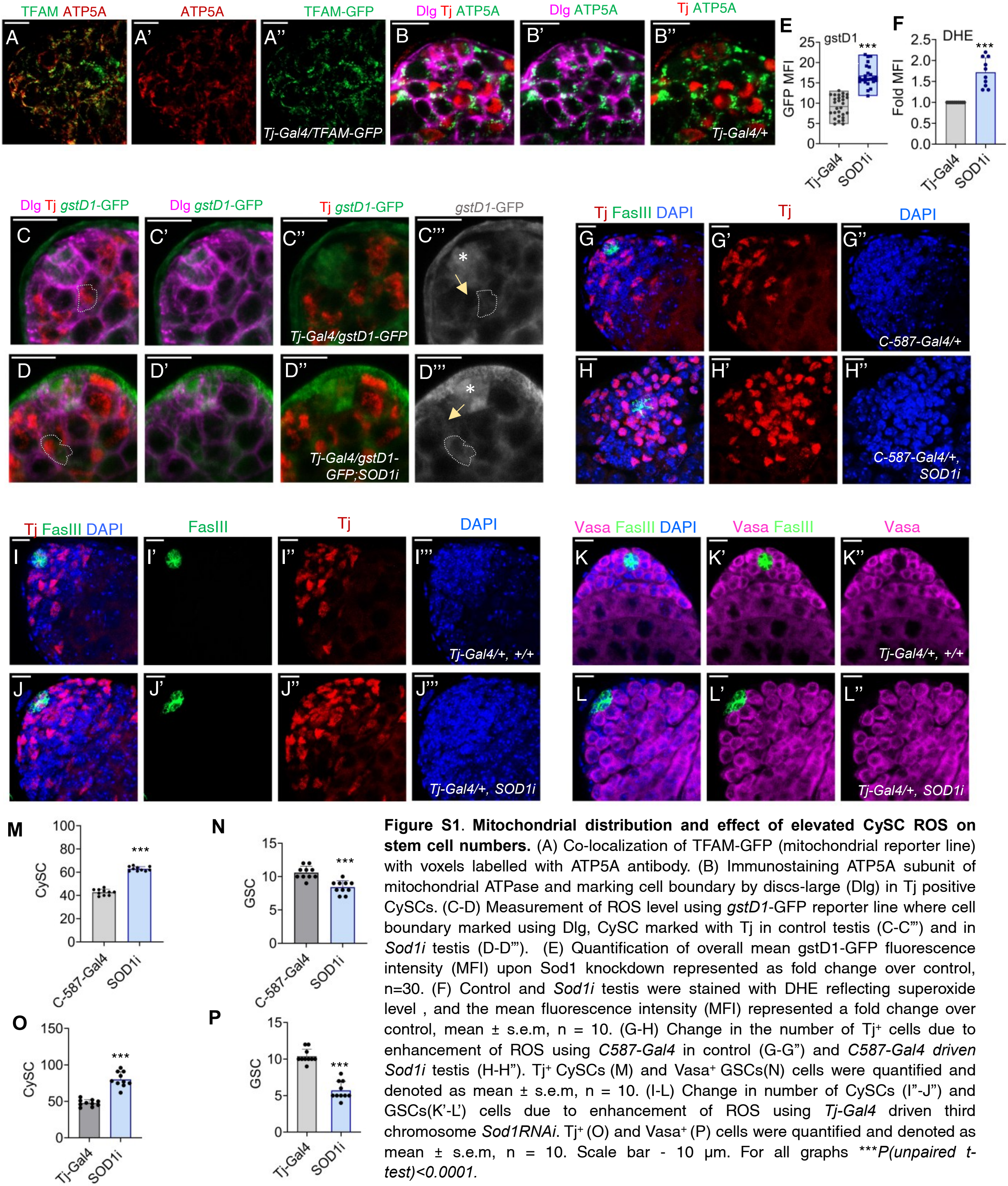


**Figure S1**. **Mitochondrial distribution and effect of elevated CySC ROS on stem cell numbers.** (A) Co-localization of TFAM-GFP (mitochondrial reporter line) with voxels labelled with ATP5A antibody. (B) Immunostaining ATP5A subunit of mitochondrial ATPase and marking cell boundary by discs-large (Dlg) in Tj positive CySCs. (C-D) Measurement of ROS level using *gstD1*-GFP reporter line where cell boundary marked using Dlg, CySC marked with Tj in control testis (C-C"') and in *Sod1i* testis (D-D"'). (E) Quantification of overall mean gstD1-GFP fluorescence intensity (MFI) upon Sod1 knockdown represented as fold change over control, n=30. (F) Control and *Sod1i* testis were stained with DHE reflecting superoxide level , and the mean fluorescence intensity (MFI) represented a fold change over control, mean ± s.e.m, n = 10. (G-H) Change in the number of Tj⁺ cells due to enhancement of ROS using *C587-Gal4* in control (G-G") and *C587-Gal4 driven Sod1i* testis (H-H"). Tj⁺ CySCs (M) and Vasa⁺ GSCs(N) cells were quantified and denoted as mean ± s.e.m, n = 10. (I-L) Change in number of CySCs (I"-J") and GSCs(K'-L') cells due to enhancement of ROS using *Tj-Gal4* driven third chromosome *Sod1RNAi*. Tj⁺ (O) and Vasa⁺ (P) cells were quantified and denoted as mean ± s.e.m, n = 10. Scale bar - 10 μm. For all graphs ****P(unpaired t-test)<0.0001.*

### Figure S2

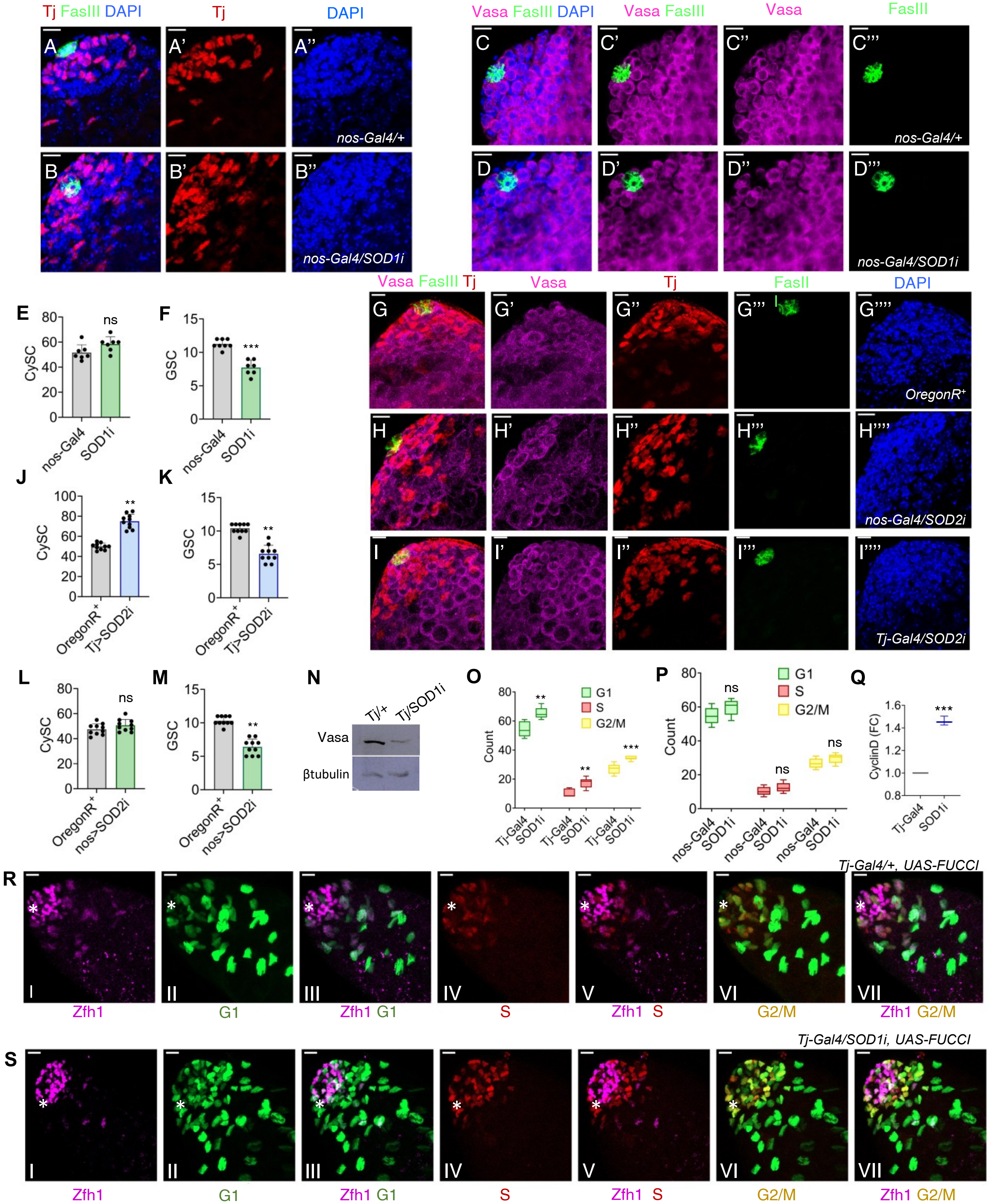

Tj FasIII DAPI
Tj
DAPI
nos-Gal4/+
nos-Gal4/SOD1i
Vasa FasIII DAPI
Vasa FasIII
Vasa
FasIII
nos-Gal4/+
nos-Gal4/SOD1i
Vasa FasIII Tj
Vasa
Tj
FasII
DAPI
OregonR+
nos-Gal4/SOD2i
Tj-Gal4/SOD2i
CySC
GSC
nos-Gal4
SOD1i
OregonR+
Tj>SOD2i
nos>SOD2i
ns
***
**
Tj/+
Tj/SOD1i
Vasa
βtubulin
Count
G1
S
G2/M
Tj-Gal4
SOD1i
CyclinD (FC)
Tj-Gal4/+, UAS-FUCCI
Tj-Gal4/SOD1i, UAS-FUCCI
Zfh1
G1
Zfh1 G1
S
Zfh1 S
G2/M
Zfh1 G2/M

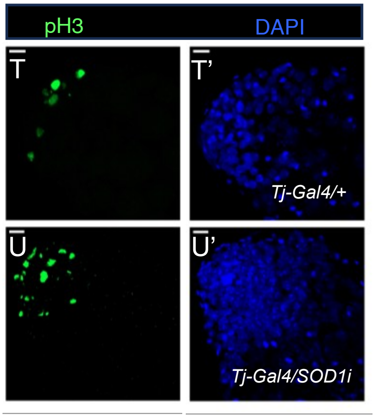


**Figure S2. Niche composition upon depletion of Sod paralogs in CySCs or GSCs.**

(A-F) Distribution of Tj$^+$ CySCs (A'-B') and Vasa$^+$ GSCs (C"-D") in control (*nos-Gal4/+*) compared to that of *nos-Gal4/SOD1i* was imaged (A-D) and quantified (E-F). (G-M) Changes in the number of Tj$^+$ (G", H" and I") and Vasa$^+$ cells (G', H' and I') when Sod2 was depleted either in CySCs (using *Tj-Gal4* driver) (I, J and K) or GSCs (using *Nos-Gal4* driver) (H, L and M). Data denotes mean ± s.e.m, n = 10. (N) Immunoblot using Vasa specific antibody showing changes in overall Vasa levels when Sod1 was knockdown. β-tubulin was used as loading control. (O-P) Assessment of cell proliferation in *Tj>Sod1i* (O) and *Nos>Sod1i* (P) using UAS-FUCCI reporter line, n=10. (Q) Quantitative RT-PCR reflecting the fold change difference of CyclinD expression in *Tj>SOD1i testes* compared to control. Data is depicted as mean ± s.e.m, n = 3. (R-S) Representative images showing variations in Zfh1$^+$ cells present in different phases of cell cycle among control (R: I-VII) and experimental flies (S: I-VII). (R:V & S:V) shows co-localization of Zfh1 with S-phase cell. Scale bar - 10 µm, ns - non-significant. Data points denotes mean ± s.e.m, n = 10. (T-U) Control (T) and *Tj-Gal4* driven *Sod1RNAi* (U) testes stained with pH3, a mitotic marker denoting the cell proliferation. For all graphs ****P(unpaired t-test)<0.0001, **P(unpaired t-test)<0.001.*

#### Figure S3

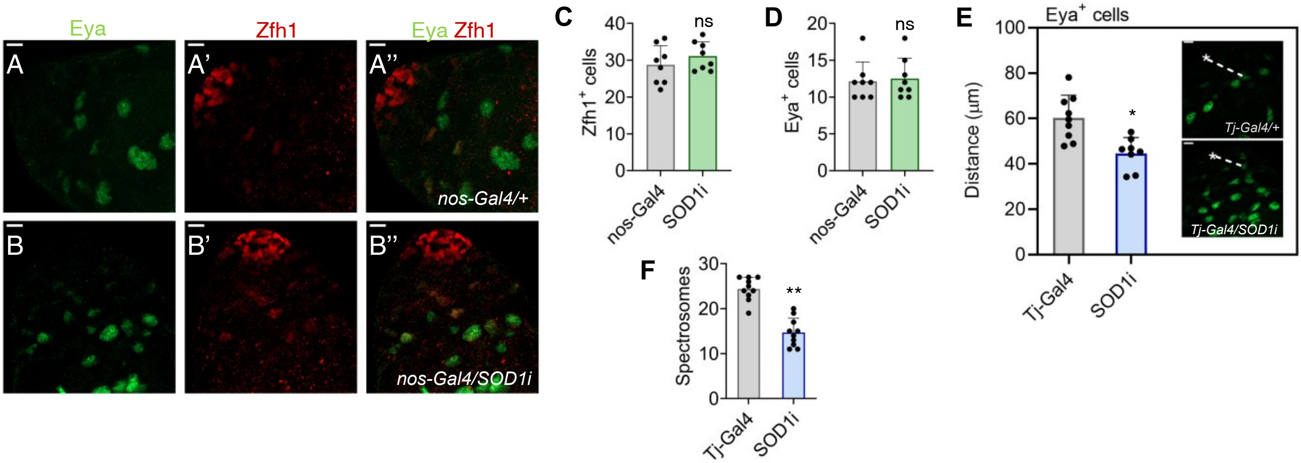


**Figure S3. Enhanced ROS in GSCs has no effect on early CySCs and late cyst cells but changes in CySCs affect differentiation and cell-cell adhesion.**

(A-D) Distribution of the Zfh1⁺ early CySCs (A'-B') and Eya⁺ late cyst cells (A-B) were imaged and quantified (C-D) under *nos>Sod1RNAi* background where (D) represents the number of Eya⁺ cells present near the niche and not the total count. (E) Mean distance between the delectable origin of anti-Eya labelled cells and hub in control and *Tj-Gal4>Sod1i* testis. (F) Total number of round spectrosome present in *Sod1i* testis with respect to controls. Data represents mean ± s.e.m, n = 10 for all the above quantification and ***P(unpaired t-test)<0.001, *P(unpaired t-test)<0.01*. ns- not significant

Figure S4

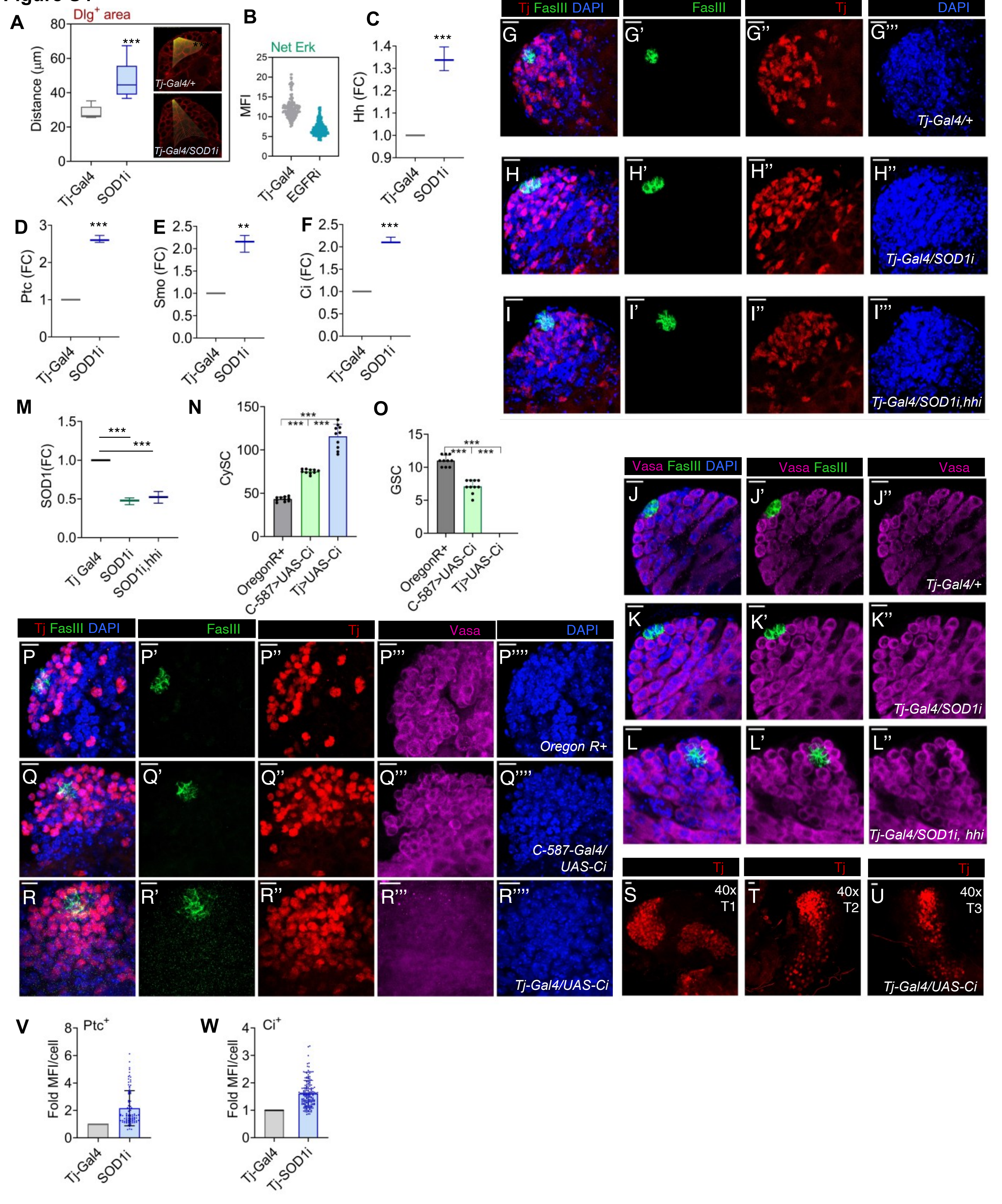

A
Dlg+ area
Distance (µm)
Tj-Gal4/+
Tj-Gal4/SOD1i
Tj-Gal4
SOD1i
***
B
Net Erk
MFI
Tj-Gal4
EGFRi
C
Hh (FC)
***
D
Ptc (FC)
E
Smo (FC)
**
F
Ci (FC)
G
Tj FasIII DAPI
FasIII
Tj
DAPI
Tj-Gal4/+
H
Tj-Gal4/SOD1i
I
Tj-Gal4/SOD1i,hhi
M
SOD1(FC)
Tj Gal4
SOD1i
SOD1i,hhi
N
CySC
OregonR+
C-587>UAS-Ci
Tj>UAS-Ci
O
GSC
J
Vasa FasIII DAPI
Vasa FasIII
Vasa
Tj-Gal4/+
K
Tj-Gal4/SOD1i
L
Tj-Gal4/SOD1i, hhi
P
Oregon R+
Q
C-587-Gal4/ UAS-Ci
R
Tj-Gal4/UAS-Ci
S
40x T1
T
40x T2
U
40x T3
Tj-Gal4/UAS-Ci
V
Ptc+
Fold MFI/cell
Tj-Gal4
SOD1i
W
Ci+
Tj-Gal4
Tj-SOD1i

**Figure S4. Activation of CySC maintenance signalling promoting CySC proliferation.** (A) Relative spread of the $Dlg^+$ cells irrespective of their expression state was calculated, n = 10. (B) MFI of total pErk expression was quantified in control and *Tj-Gal4* driven *EGFR RNAi,* n=200 (C-F) Quantitative RT-PCR reflecting the fold changes differences among Hh (C), Ptc (D), Smo (E) and Ci (F) expression in *Tj>SOD1i testes* compared to control. Data is depicted as mean ± s.e.m, n = 3. (G-L) Representative images showing distribution of Tj (G'',H'', and I'') and Vasa (J'', K'' and L'') labelled cells in control, *Tj-Gal4* driven *Sod1i*, and *Sod1i/Hhi* .(M) Quantitative RT-PCR reflecting the fold changes differences of SOD1 level among control, single UAS and double UAS construct. Data is depicted as mean ± s.e.m, n = 3. (N-R) Distribution of Tj (P'',Q'', and R'') and Vasa (P''',Q''', and R''') labelled cells were imaged in control (P-P''''), *C-587 Gal4* driven *UAS-Ci* (Q-Q''') and *Tj-Gal4* driven *UAS-Ci* (R-R''') and quantification of CySC number (N) and GSC number (O). (S-U) More representative images of *Tj-Gal4* driven *UAS-Ci* under 40x magnification showing over-proliferative Tj+ cells. (V, W) Fold Ptc and Ci levels with respect controls, normalized using total $Tj^+$ cells near the hub, n (Ptc) = 90, n (Ci) = 170, ****P(unpaired t-test)<0.0001.* Scale bar - 10 μm.

#### Figure S5

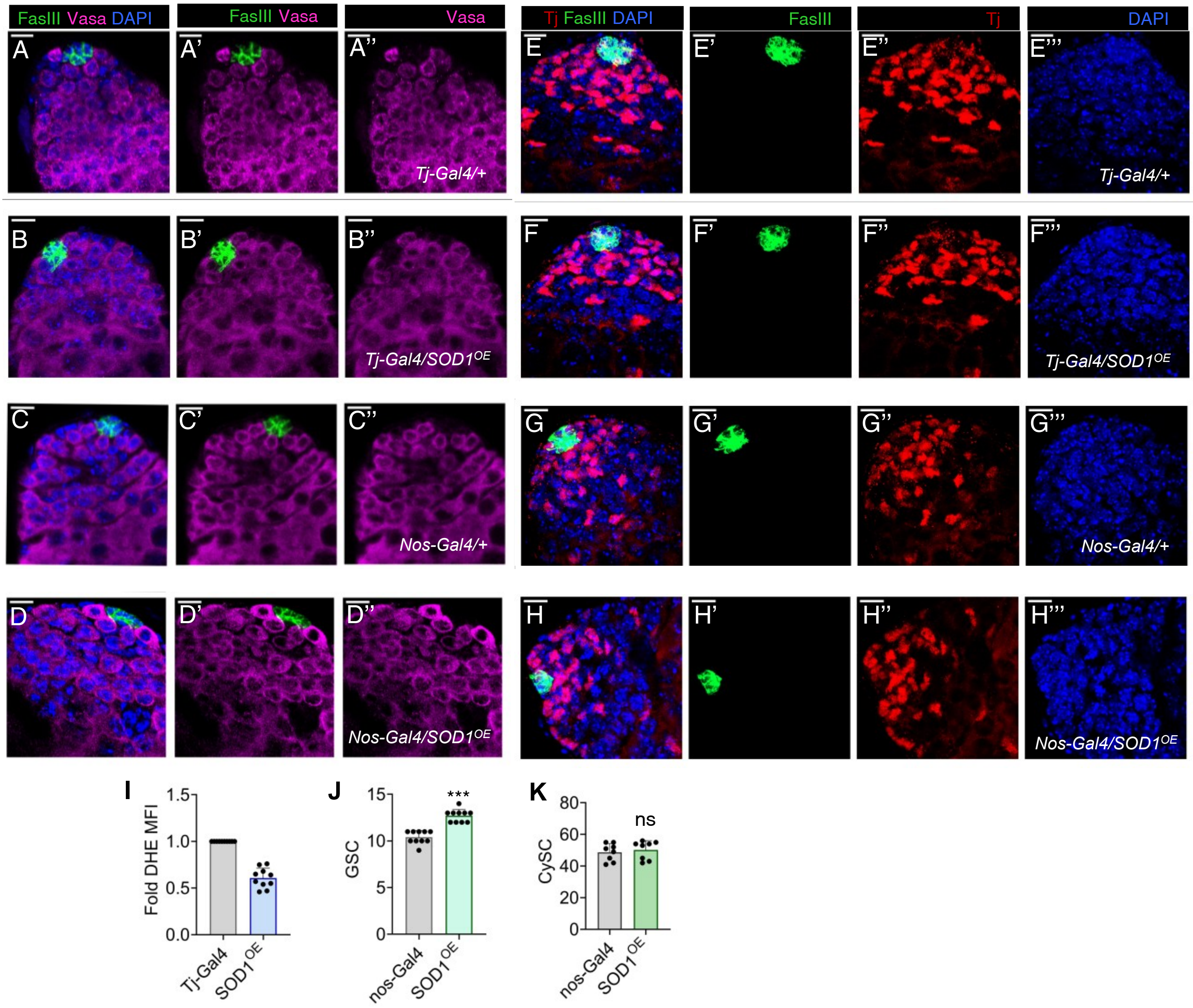


**Figure S5. Overexpression of Sod1 either in GSCs or CySCs sustains self-renewing propensity of GSC-like cells.**

(A''-D'') Images showing rosette arrangement of GSCs around the hub in control and experimental testis overexpressing Sod1 in CySCs (*Tj-Gal4/Sod1$^{OE}$*) and GSCs (*nos-Gal4/Sod1$^{OE}$*). (E-H) The number of Tj$^{+}$ cells in *Tj-Gal4/Sod1$^{OE}$* and *nos/Sod1$^{OE}$* testis were imaged (E''-H''). (I) Fold DHE mean fluorescence intensity of *Tj-Gal4/Sod1$^{OE}$* testis over controls. The GSCs (J) and CySCs (K) were quantified over control and represented as bar graph. Data denotes mean ± s.e.m., n = 10, ****P(unpaired t-test)<0.0001, ns – not significant.*
